# Cell Type-Specific Modulation of Layer 6A Excitatory Microcircuits by Acetylcholine in Rat Barrel Cortex

**DOI:** 10.1101/701318

**Authors:** Danqing Yang, Robert Günter, Guanxiao Qi, Gabriele Radnikow, Dirk Feldmeyer

**Author notes:** Corresponding author: Dirk Feldmeyer, Institute of Neuroscience and Medicine (INM-10) Research Centre Jülich D-52425 Jülich Germany.

## Abstract

Acetylcholine (ACh) is known to regulate cortical activity during different behavioral states, e.g. wakefulness and attention. Here we show a differential expression of muscarinic ACh receptors (mAChRs) and nicotinic AChRs (nAChRs) in different layer 6A (L6A) pyramidal cell (PC) types of somatosensory cortex. At low concentrations, ACh induced a persistent hyperpolarization in corticocortical (CC) but a depolarization in corticothalamic (CT) L6A PCs via M_4_ and M_1_ mAChRs, respectively. At ∼1 mM ACh depolarized exclusively CT PCs via α_4_β_2_ subunit-containing nAChRs without affecting CC PCs. Miniature EPSC frequency in CC PCs was decreased by ACh but increased in CT PCs. In synaptic connections with a presynaptic CC PC, glutamate release was suppressed via M_4_ mAChR activation but enhanced by nAChRs via α_4_β_2_ nAChRs when the presynaptic neuron was a CT PC. Thus, in layer 6A the interaction of mAChRs and nAChRs results in an altered excitability and synaptic release, effectively strengthening corticothalamic output while weakening corticocortical synaptic signaling.

## Introduction

Acetylcholine (ACh) has been shown to play a major role in memory processing, arousal, attention and sensory signaling^1-7^. It has been demonstrated that the ACh concentration in the cerebrospinal fluid increases during wakefulness and sustained attention^8,9^. In the neocortex release of ACh occurs predominately via afferents originating from cholinergic neurons in the nucleus basalis of Meynert of the basal forebrain^10-12^; their terminals are densely distributed throughout all neocortical layers^13-15^. A classical view is that ACh invariably increases the excitability of excitatory neurons in neocortex 16-20. However, a persistent hyperpolarization in layer 4 (L4) excitatory neurons was found in somatosensory cortex^21,22^. This layer-specific cholinergic modulation may contribute to improving the cortical signal-to-noise ratio^23-25^.

Although extensive studies have been conducted on the cholinergic modulation of neocortical excitatory neurons, the action of ACh on the layer 6 (L6) microcircuitry has not been systematically investigated. Two main pyramidal cell (PC) classes exist in cortical layer 6, namely corticothalamic (CT) and corticocortical (CC) PCs. These two neuron types differ in their axonal projection patterns, dendritic morphological features, electrophysiological properties and expression of molecular markers^26-30^. CC PCs have no subcortical target and send intracortical projections mainly within the infra-granular layers^28^; CT PCs, in contrast, have few axons distributed in cortex and send projections directly back to the thalamus thereby contributing to a feedback control of sensory input^31-35^. The question how the function of these two classes of L6 PCs is modulated by ACh has so far not been explored.

Recent optogenetic studies suggest that PCs in L5 and L6 receive direct cholinergic inputs^20,36^. In these neurons, ACh induces a slowly desensitizing inward current in L6 PCs of prefrontal cortex through activation of α_4_β_2_ subunit containing synaptic nicotinic acetylcholine receptors (nAChRs)^23,36-39^. However, there are very few studies focusing on the effects of muscarinic acetylcholine receptors (mAChRs) in L6A neurons^29,40^. Here, using single and paired patch-clamp recordings with simultaneous biocytin-filling, we investigated both muscarinic and nicotinic modulation of morphologically identified excitatory neurons and their synaptic connections in layer 6A of rat primary somatosensory barrel cortex. We found that ACh shows a cell type-specific effect on both cellular and synaptic properties in L6A excitatory microcircuits through activation of mAChRs and/or nAChRs. Our results reveal that two functionally and morphologically distinct subpopulations of L6A PCs, CC and CT PCs, are differentially modulated by ACh. We demonstrate that ACh suppresses intracortical synaptic transmission via somatodendritic hyperpolarization and inhibition of presynaptic neurotransmitter release of CC PCs by activating M_4_Rs. In contrast, CT PC show a dual cholinergic modulation: These neurons are depolarized via M_1_ mAChRs and α_4_β_2_ subunit-containing nAChRs while the presynaptic release probability is enhanced by α_4_β_2_ nAChRs. In this way, ACh contributes to a facilitation of corticothalamic feedback.

## Results

### ACh either depolarizes or hyperpolarizes L6A PCs through activation of mAChRs

Whole-cell patch clamp recordings from L6A neurons were performed in acute brain slices of rat barrel cortex with simultaneous biocytin fillings. During recordings, excitatory neurons were distinguished from interneurons by their regular firing pattern with a low maximum firing frequency. Following bath application of 100 µM ACh, one subset of L6A PCs showed a membrane potential hyperpolarization by on average -2.0 ± 1.0 mV (n = 14), whereas another was depolarized by +9.5 ± 6.1 mV (n = 15; **Fig. 1a**). In addition, 1 s current pulses were injected in the recorded neuron to elicit AP firing before and during bath application of ACh. Under the supra-threshold stimulus (100pA above the rheobase current), the firing frequency was decreased by ACh in L6A PCs showing an ACh-induced hyperpolarization but increased in PCs that exhibit a depolarizing ACh response (**Fig. 1a, b**). Notably, both ACh-induced hyperpolarization and depolarization were not transient but persisted until the end of bath application. For L2/3 and L5 PCs it has been reported that the ACh-induced depolarization was preceeded by an initial transient hyperpolarization mediated by ‘small conductance’, Ca^2+^-activated K^+^ channels^21,41-43^. We were able to reproduce this finding under the same recording condition; however, the de- and hyperpolarizing cholinergic response in L6A PCs induced by ACh puff application was always monophasic (**Supplementary Fig. 1**).

**Fig. 1.**
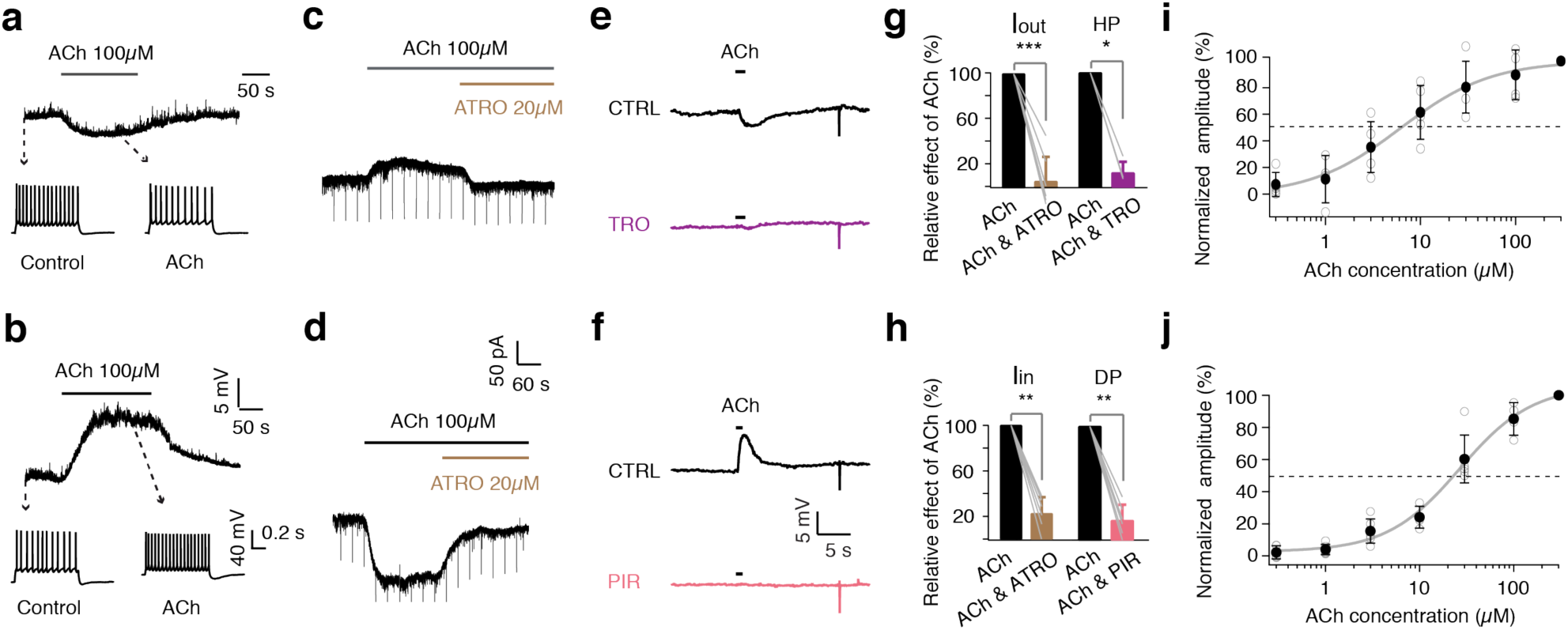
Low concentrations of ACh induce either a hyperpolarization or a depolarization of L6A PCs by activating mAChRs. **(a**, **b)** Top, L6A PC either show a hyperpolarizing (a) or a depolarizing (b) response following bath application of 100 µM ACh. Bottom, firing patterns of neurons in response to 1 s depolarising current injection (rheobase + 100 pA) before and during ACh application. **(c**, **d)** Representative voltage-clamp recordings with bath application of ACh showing outward current (I_out_) (e) or inward current (I_in_) (f) in L6A PCs. The effects are blocked by 20 µM atropine (ATRO). **(e**, **f)** Representative current-clamp recordings showing that puff application of ACh (100µM) evokes a fast hyperpolarization (HP) (g) or depolarization (DP) (h) in L6A PCs. The specific M_1_ mAChR antagonist pirenzipine (PIR, 0.5 µM) or the specific M_4_ mAChR antagonist tropicamide (TRO, 1 µM), respectively, were added to the perfusion solution to block ACh-induced membrane potential changes. **(g**, **h)** Summary bar graphs showing the percentage block by general and specific mAChR antagonists. n = 7, p = 0.0006 for I_out_ group, n = 7, p = 0.0022 for I_in_ group, n = 4, p = 0.029 for HP group and n = 6, p = 0.0022 for DP group. Statistical analysis was performed using Mann Whitney *U*-test. Error bars represent SD. **(i**, **j)** Muscarinic responses of ACh were examined in the presence of 1 µM mecamylamine (MEC) and 0.5 µM TTX. ACh dose-response curves for hyperpolarizing (n = 5) (c) and depolarizing (n = 8) (d) L6A PCs are fitted by the Hill equation. Dashed lines represent half maximal effects. The corresponding EC_50_ is 6.2 ± 1.3 µM for hyperpolarizing PCs and 26.7 ± 5.4 µM for depolarizing PCs. Filled circles represent mean values of different ACh concentrations.

To determine which fraction of the membrane potential changes in L6A pyramidal neurons is mediated by mAChRs, 20 µM atropine (ATRO, a general mAChR antagonist) was applied in voltage-clamp mode. Both the ACh-induced outward and inward currents were found to be strongly blocked by ATRO (20 µM) **(Fig. 1 c, d)**, suggesting that both ACh response types in L6A excitatory neurons are almost exclusively mediated by mAChRs. We hypothesized that the G_i/o_ protein coupled M_4_ mAChR subtype (M_4_Rs) mediates the hyperpolarizing effects while the G_q/11_ proteincoupled M_1_ mAChR subtype (M_1_Rs) is responsible for the depolarization induced by ACh application. To test this, puff application of ACh (100 µM) was performed in the presence and absence of the selective mAChR antagonists in the perfusion solution. In the presence of 1 µM tropicamide (TRO, a selective M_4_R antagonist), the ACh-induced hyperpolarization was abolished (**Fig. 1e, g**). Conversely, the ACh-induced depolarization was blocked by 0.5 µM pirenzipine (PIR, a selective M_1_R antagonist; **Fig. 1f, h**). These results indicate that the persistent hyperpolarization and depolarization induced by low concentrations of ACh are mediated exclusively by M_4_Rs and M_1_Rs, respectively.

The dose-dependence of the muscarinic effects was investigated by bath application of increasing concentrations of ACh in the presence of 1 µM mecamylamine (MEC, a general nAChR antagonist) and 0.5 µM tetrodotoxin (TTX) (0.3 µM to 300 µM; **Fig. 1i, j**). The dose-response curve was obtained by fitting the data to the Hill equation. For hyperpolarizing L6A PCs, the ACh concentration for a half-maximum response (EC_50_) was 6.2 ± 1.3 µM while for depolarizing neurons, the EC_50_ was 26.7 ± 5.4 µM. Thus, an ACh concentration of 30 µM was adopted for all subsequent experiments; this concentration resulted in a >50% of the maximum response in both subgroups of L6A excitatory neurons. In addition, when only 30 µM ACh was used neurons did not respond with AP firing which was occasionally observed when applying 100 µM ACh.

### Cholinergic responses in L6A PCs are cell-type specific

To investigate whether the two different cholinergic response types are specific for a defined L6A pyramidal cell type, we characterized L6A PCs by their morphological, electrophysiological and molecular features. Here, a total of 105 excitatory L6A neurons were recorded and morphologically reconstructed. Previous studies have consistently shown that CC and CT L6A PCs can be distinguished reliably by their axonal projection patterns^26,27,30,44^. Of all excitatory cells, 74 (70.5%) were identified as putative CT PCs while 31 (29.5%) were putative CC PCs. CC L6A PCs displayed a dense horizontal axonal projection pattern in infragranular layers spanning several neighboring barrel columns; CT L6A PCs, on the other hand, showed a sparse columnar axonal domain with the majority of collaterals projecting directly towards the pia and terminating predominately in layer 4 **(cf Fig. 2a left and right panels**). CC PCs have a significantly larger axonal (15523 ± 5013 µm vs. 5209 ± 1462 µm, P < 0.001) and dendritic length (5921 ± 1346 µm vs. 5134 ± 1070 µm, P < 0.05) compared to CT PCs. Similar differences were also detected in the horizontal axonal and dendritic field span (1714 ± 350 µm vs. 358 ± 111 µm, P < 0.001 and 361 ± 58 µm vs. 232 ± 28 µm, P < 0.001, respectively, for CC vs. CT L6A PCs). For CC L6A PCs these values are likely to be strong underestimates (by ≥ 90%, cf.^45^) because in acute slice preparations their long-range axonal collaterals will be severely truncated; however, this does not prevent an unambiguous cell type identification. In addition, CC L6A PCs have more first order axon collaterals (p < 0.001) but fewer dendrites (p < 0.001) than CT PCs **(Fig. 2b)**. The features described above are reflected in the polar plots **(Fig. 2a)**.

**Fig. 2.**
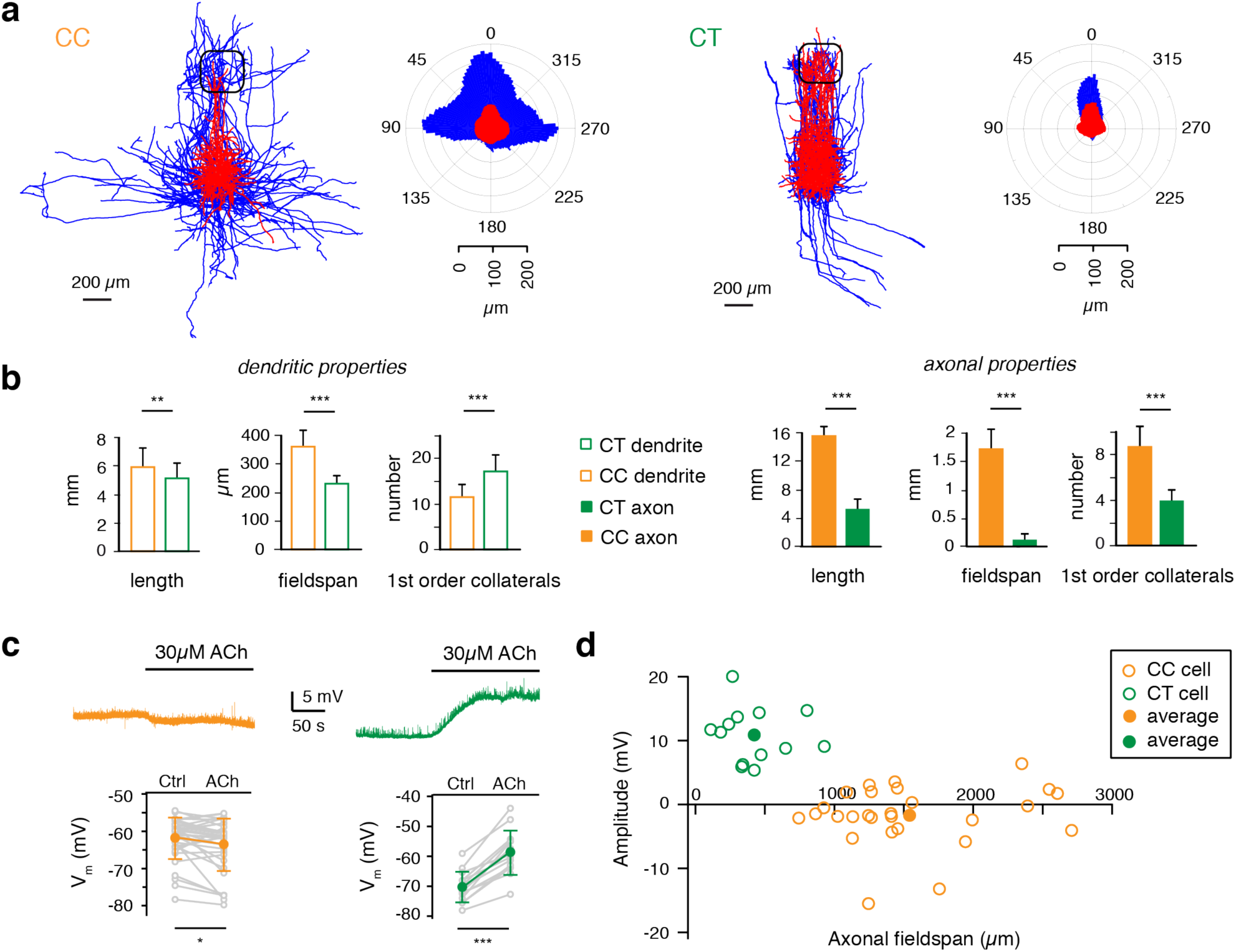
ACh hyperpolarizes CC L6A PCs but depolarizes CT L6A PCs. **(a)** Left, overlay of reconstructions of CC and CT PCs. Reconstructions of PCs were aligned with respect to the barrel center. Right, polar plots of CC and CT PCs. n = 15 for each group. Somatodendrites are shown in red and axons are shown in blue. **(b)** Histograms comparing the length, fieldspan and number of first order collaterals of axonal and dendritic structures for the two groups of PCs. n = 21 for CC neurons and n = 54 for CT neurons. Dendritic length: p = 0.0015, dendritic fieldspan: p = 1.4 × 10^-10^, number of dendritic main nodes: p = 1.7 × 10^-6^; Axonal length: p = 9.6 × 10^-8^, dendritic fieldspan: p = 4.8 × 10^-11^, number of axoanl main nodes: p = 8.5 × 10^-11^ for Mann-Whitney *U*-test. **(c)** Top, representative current-clamp recordings of a depolarizing CC (orange) and a hyperpolarizing CT pyramidal cell (green) following bath application of 30µM ACh. Bottom, histograms of resting membrane potential (V_m_) of L6A CC PCs in control and in the presence of 30 µM ACh (n = 35, p = 0.019 for Wilcoxon signed-rank test) and CT (n = 14, p = 6.1 × 10^-5^ for Wilcoxon signed-rank test) PCs. **(d)** Plots of the ACh-induced change in V_m_ vs axonal fieldspan for two subtypes of PCs. Open orange circles, data from individual CC PCs (n = 27); open green circles, data from individual CT PCs (n = 13). Filled orange circle, average data from CC cells; filled green circle, average data from CT cells.

In addition, we determined the electrophysiological properties of morphologically identified CC (n = 11) and CT (n = 9) L6A PCs. Compared to CT PCs, CC PCs showed a significantly lower Rin (P < 0.05), a longer onset time (P < 0.01) for the first action potential (AP) evoked by injecting a rheobase current and a longer AP half-width (P < 0.05). Trains of spikes were elicited to examine the firing behavior. The AP adaptation ratio (2^nd^ ISI/10^th^ ISI) of CC PCs was smaller (P < 0.05) than that of CT cells because they exhibited an initial spike burst. **(Supplementary Fig. 2)**. The differences in passive and active electrophysiological properties found here are in accordance with previous studies^26,40^.

Furthermore, the nuclear transcription factor Fork-head box protein P2 (FoxP2) is co-expressed with the neurotensin receptor 1 (NtsR1) gene, a molecular marker for CT L6A PCs in mice^29,46^. To identify the expression of FoxP2 in L6A PCs, we performed whole-cell recordings with simultaneous filling of biocytin and fluorescent Alexa Fluor® 594 dye (n = 14). Subsequently, brain slices were processed for FoxP2 immunofluorescence staining. We found that CT L6A PCs were FoxP2-positive while CC PCs are FoxP2-negative **(Supplementary Fig. 3a, b)**. The tight correlation between neuronal morphology, electrophysiology and FoxP2 expression demonstrates the reliability of classification based on axonal projection patterns of CC and CT PCs.

ACh at a concentration of 30 µM was bath-applied to 63 morphological identified L6A neurons. CC L6A PCs showed a hyperpolarizing response with a mean amplitude of -1.76 ± 4.28 mV (from -61.9 ± 5.6 mV to -63.6 ± 7.0 mV, P < 0.05, n = 35). In contrast, ACh (30 µM) induced a strong depolarization with a mean amplitude of +11.4 ± 4.6 mV (from -70.3 ± 5.1 mV to -58.8 ± 7.4 mV, P < 0.001, n = 14) in CT PCs without exception (**Fig. 2c)**. In **Fig. 2d**, ACh-induced membrane potential changes are plotted against the horizontal axonal field span revealing a strong correlation between axonal morphology and cholinergic response for the two L6A pyramidal cell types. In addition, by performing immunostaining we confirmed that M_4_Rs were enriched within L6A. We found that M_4_R-positive neurons were FoxP2-negative while virtually no FoxP2-positive neuron expressed M_4_Rs **(Supplementary Fig. 3d)**. This is consistent with our pharmacological result that only FoxP2-negative CC PCs showed a M_4_Rs-mediated hyperpolarization following ACh application **(Supplementary Fig. 3)**.

### CT PCs are selectively activated by high concentrations of ACh via α4β2 nAChRs

As demonstrated above, the depolarizing and hyperpolarizing effects of ACh in L6A PCs can be attributed to the activation of M_1_ and M_4_ mAChRs, respectively (**Fig. 1c, d**). However, previous studies have shown that ACh excites L6A excitatory neurons by activating nAChRs^23,36,37,39^. In order to investigate the functional role of nAChRs in L6A of rat barrel cortex, we perfused slices continuously with 200 nM ATRO. Under this condition, 30 µM ACh had no effect on both CC and CT PCs (P > 0.05 for CC cells, n = 5; P > 0.05 for CT cells, n = 5) (**Fig. 3a**); application of 1 mM ACh, however, strongly depolarized CT PCs (P < 0.001, n = 12) while CC PCs showed no response (P > 0.05, n = 4; **Fig. 3b**). Our results demonstrate that both the muscarinic and nicotinic modulation of L6A PCs is cell-type specific; nAChRs are present solely in CT L6A PCs and activated substantially only by high ACh concentrations. To determine the concentration range in which ACh activates postsynaptic nAChRs, we measured the dose-response curve for ACh in the presence of 200 nM ATRO. A fit of dose-response relationship to the Hill equation gave an EC_50_ of 1.2 ± 0.3 mM (n = 5) for the nicotinic ACh response (**Fig. 3c**), a value more than about two orders of magnitude larger than those of the de- and hyperpolarizing muscarinic response.

It has been reported that the expression of nAChR subtypes in the neocortex exhibits layer-specificity. L6A PCs in prefrontal cortex show a slow inward current to ACh by activating nAChRs containing the α_4_ and β_2_ subunits^23,36^. To confirm that this nAChR subtype mediates the response, the response of CT L6A PCs to application of 1 mM ACh was recorded in an ATRO-containing perfusion solution. In the presence of DHßE (10 µM), a nicotinic antagonist specific for α_4_β_2_ nAChRs, the ACh-dependent depolarization in CT PCs was eliminated, suggesting that CT L6A PCs express postsynaptic α_4_β_2_ subunit-containing nAChRs (**Fig. 3d**).

**Fig. 3.**
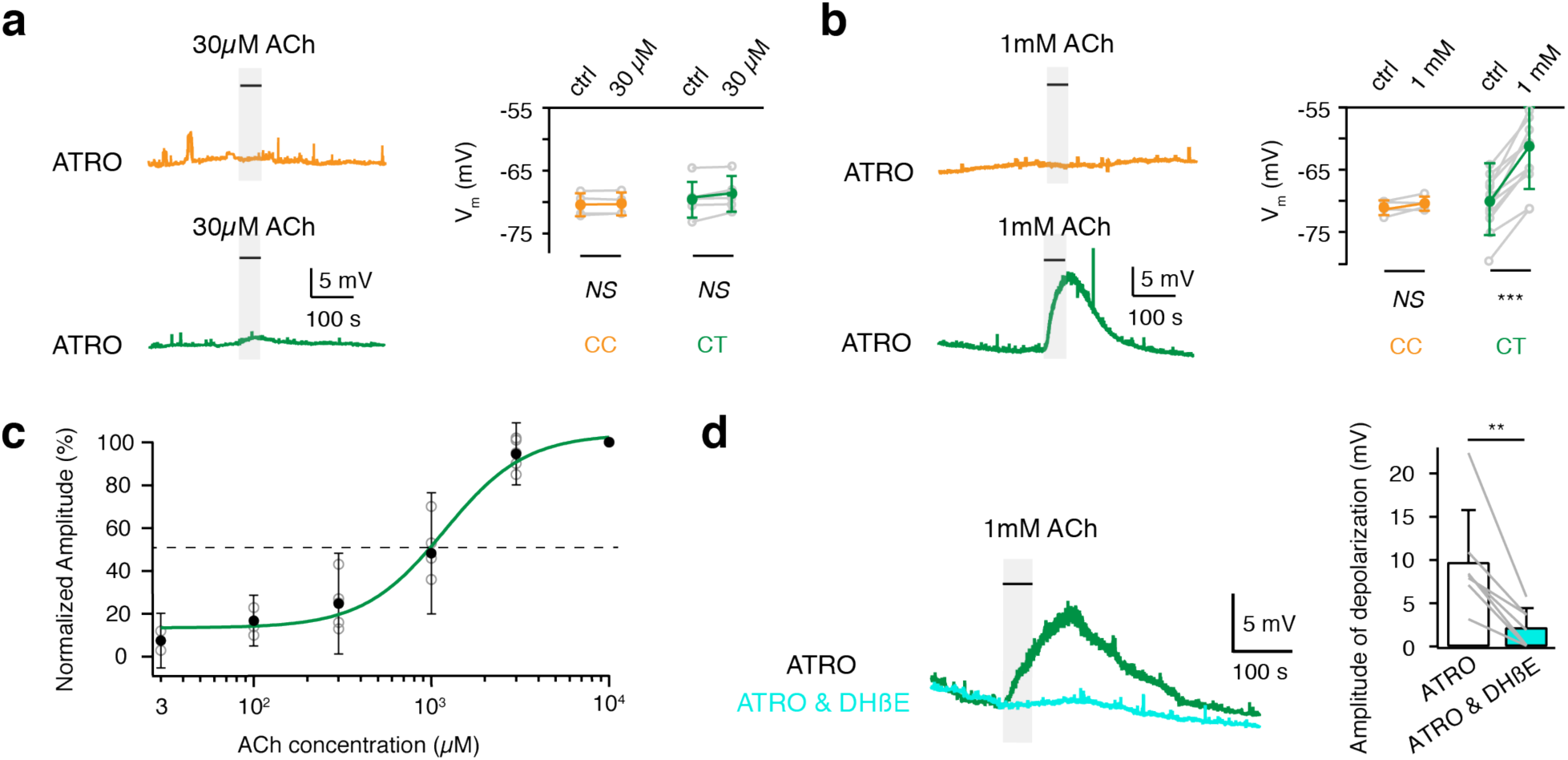
High concentration of ACh selectively depolarizes CT PCs. **(a)** In the presence of 200 nM atropine (ATRO), bath application of low concentration ACh (30 µM, 50s) show no effect on either CC (top) or CT (bottom) PCs. Summary plots show the resting membrane potential (V_m_) under control and ACh conditions in L6A CC (n = 5, p = 0.188) and CT (n = 5, p = 0.0625) PCs. NS (not significant) for Wilcoxon signed-rank test. Error bars represent SD. **(b)** In the presence of 200 nM ATRO, bath application of 1 mM ACh for 50s has no effect on CC L6A PCs (top) but induces a strong depolarization of CT L6A PCs (bottom). Summary plots showing V_m_ of CC L6A PCs under control conditions and in the presence of ACh in L6A CC (n = 5, p = 0.156) and CT (n = 12, p = 0.0002) PCs. Statistical analysis was peformed using Wilcoxon signed-rank test. Error bars represent SD. **(c)** The dose-response curve of ACh under ATRO application (200 nM) in CT PCs (n = 5) is well fitted by the Hill equation. The dashed line indicates the half maximal effect; the corresponding EC_50_ is 1.2 mM. Filled circles show mean effect of different concentrations while open circles represent individual values. Error bars represent SD. **(d)** The depolarization induced by ACh application (in the presence of 200 nM ATRO) is blocked by 10 µM of the specific antagonist of α_4_β_2_ subunit-containing nAChRs DHßE in CT L6A PCs. Summary plots showing the amplitude of the depolarization in response to application of 1 mM ACh in the presence of ATRO alone (open bar) and ATRO together with DHßE (n = 7, p = 0.0078 for Wilcoxon signed-rank test). Error bars represent SD.

### ACh differentially modulate miniature spontaneous activity of CC and CT L6A PCs

In addition to changing the membrane properties and excitability of neurons, ACh is also a powerful modulator of neurotransmitter release. Therefore, we measured the amplitude and frequency of miniature spontaneous activity by performing whole-cell voltage-clamp recordings from L6A PCs. The membrane potential was held at -70 mV and inward miniature EPSCs (mEPSCs) were recorded in the presence of TTX (0.5 µM) and gabazine (10 µM).

We found that ACh differentially modulates miniature spontaneous activity in both L6A CC and CT PCs. The frequency but not the amplitude of mEPSCs in CC L6A PCs was significantly decreased by application of 30 µM ACh (2.8 ± 0.8 vs. 2.2 ± 1.0 Hz; n = 7, P < 0.01), an effect that was blocked by the M_4_Rs antagonist tropicamide (2.2 ± 1.0 vs. 2.8 ± 1.0 Hz; n = 7, P < 0.01) (**Fig. 4a-c**). This suggests that ACh decreases the neurotransmitter release probability at synapses with CC L6A PCs via presynaptic M_4_Rs. Similarly, when DHßE was co-applied with tropicamide and ACh, a reduction of mEPSC frequency without a change in mEPSC amplitude was observed (**Supplementary Fig. 4**). This implies that in addition to M_4_Rs, α_4_β_2_ nAChRs also play a role in the cholinergic modulation of excitatory synaptic transmission onto CC PCs.

**Fig. 4.**
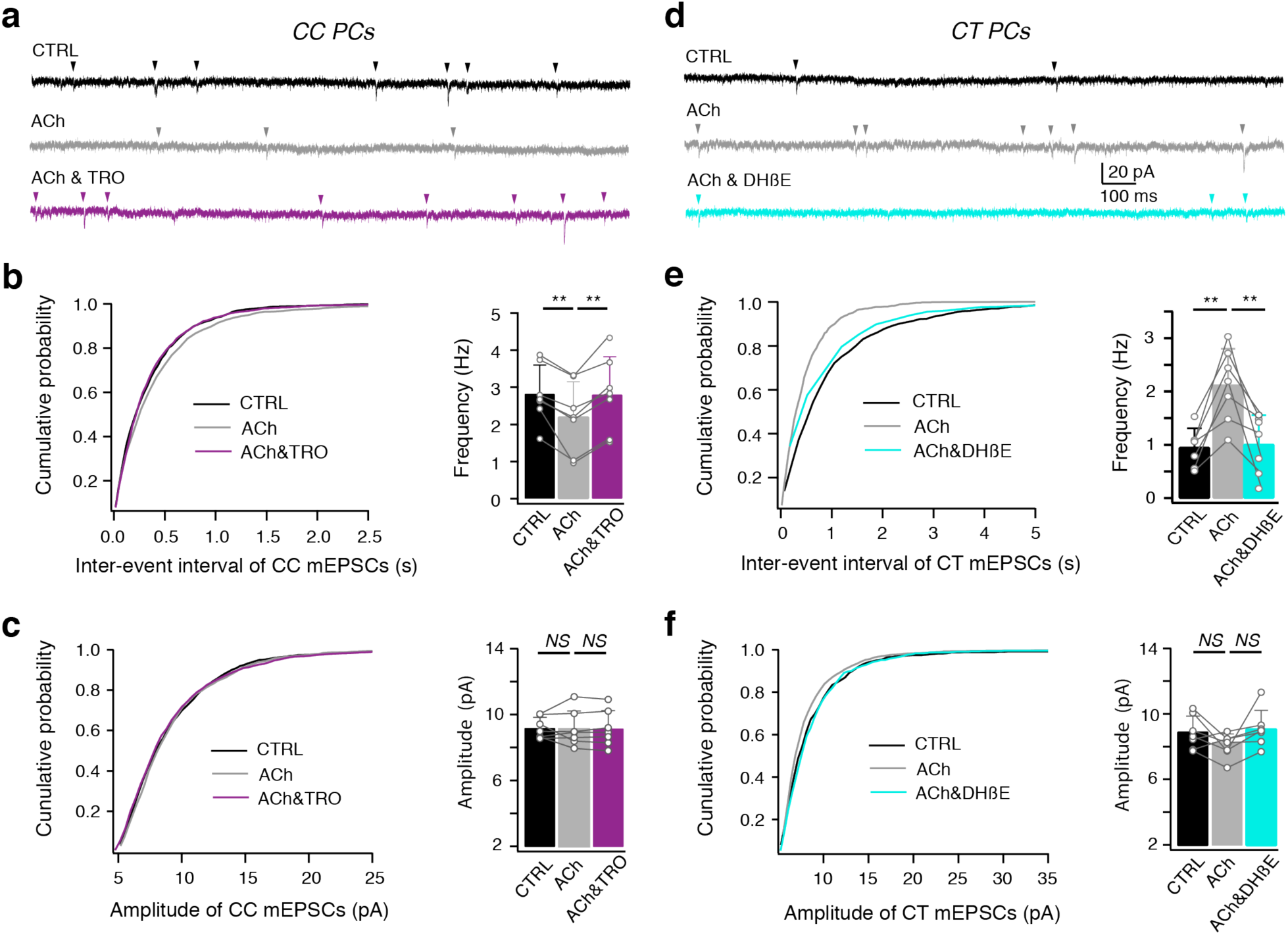
ACh differentially modulates miniature spontaneous activity in CC and CT L6A PCs. **(a)** Example voltage-clamp recordings of a CC L6A PCs under control (black), bath application of 30 µM ACh (gray) and co-application of ACh and 1 µM TRO (purple). Miniature EPSCs were recorded in the presence of TTX (0.5 µM) and GABAzine (10 µM) at a holding potential of -70 mV. **(b)** Cumulative distributions of mEPSCs inter-event interval recorded in CC L6A PCs under control condition, in the presence of ACh alone, and of ACh & TRO. Summary histograms of mEPSC frequency are shown on the right. **P < 0.01 for Wilcoxon signed-rank test. Error bars represent SD. **(c)** Cumulative distributions of mEPSCs amplitude recorded in CC L6A PCs under control, ACh and ACh & TRO conditions. Summary histograms of mEPSC amplitude are shown on the right. Control vs. ACh, p = 0.8125; ACh vs. ACh & TRO, p = 0.9375, n = 7 for Wilcoxon signed-rank test. Error bars represent SD. **(d)** Example voltage-clamp recordings of a CT L6A PC in control (black), after bath application of 30 µM ACh (gray) and subsequent co-application of ACh and 10 µM DHßE (turquoise). Miniature EPSCs were recorded in the presence of TTX (0.5 µM) and GABAzine (10 µM) at a holding potential of -70 mV. **(e)** Cumulative distributions of mEPSCs inter-event interval recorded in CT L6A PCs under control, ACh and ACh & DHßE conditions. Summary histograms of mEPSC frequency are shown on the right. **P < 0.01 for Wilcoxon signed-rank test. Error bars represent SD. **(f)** Cumulative distributions of mEPSCs amplitude recorded in CT L6A PCs under control, ACh and ACh & DHßE conditions. Summary histograms of mEPSC amplitude are shown on the right. Control vs. ACh, p = 0.4258; ACh vs. ACh & DHßE, p = 0.0781, n = 7 for Wilcoxon signed-rank test. Error bars represent SD.

In contrast to CC L6A PCs, application of 30 µM ACh significantly decreased the inter-event interval of mEPSCs in CT PCs, reflecting an increase in mEPSC frequency (0.95 ± 0.36 vs. 2.12 ± 0.68 Hz; n = 7, P < 0.01) while the mEPSc amplitude remained unaffected. Because ATRO did not affect the mEPSC frequency, we argued that the cholinergic effects on spontaneous mEPSCs in CT L6A PCs were not mediated by mAChRs but exclusively by presynaptic nAChRs. To test this, 10 µM DHßE was co-applied with ACh. In the presence of DHßE, the ACh-induced increase of mEPSCs frequency in CT L6A PCs was reduced to control level (2.12 ± 0.68 vs. 1.01 ± 0.55; n = 7, P < 0.05; **Fig. 4d-f**). These results suggest that ACh potentiates excitatory synaptic transmission onto L6A CT PCs exclusively via presynaptic α_4_β_2_ subunit containing nAChRs.

### ACh induces a reduction of presynaptic neurotransmitter release in CC L6A PCs but an an increase in CT PCs

To elucidate cholinergic effects on L6A PCs at pre- and postsynaptic sites independently, paired recordings and simultaneous biocytin fillings of synaptically coupled L6A neurons were performed. Excitatory neurons were classified as either CT and CC PCs based on the criteria mentioned above. During recordings, inhibitory interneurons were preliminarily identified by their high frequency AP firing pattern. After reconstructions, interneurons were further distinguished based on morphological features such as lack of dendritic spines. 34 excitatory connections were established by presynaptic CC PCs. We found that ACh suppresses the synaptic efficacy of neuronal connections established by presynaptic CC PCs regardless of the postsynaptic neuron type **(Fig. 5)**. The unitary EPSP (uEPSP) amplitude of all synaptic connections with a presynaptic L6A CC PC were all significantly decreased by ACh (30 µM). For CC-CC connections the uEPSP amplitude decreased from 0.45 ± 0.32 mV to 0.19 ± 0.14 mV (n = 20 pairs, P < 0.001) and for CC-CT connections changed from 0.35 ± 0.22 mV to 0.19 ± 0.13 mV (n = 5 pairs, P < 0.05). For CC-interneuron connections the mean uEPSP was reduced from 0.90 ± 0.90 mV to 0.52 ± 0.58 mV (n = 9 pairs, P < 0.05) in the presence of ACh. ACh also significantly increased the paired-pulse ratio (PPR) of CC-CC (1.0 ± 0.4 vs. 1.5 ± 0.7, P < 0.01), CC-CT (1.2 ± 0.7 vs. 2.1 ± 1.6, P < 0.05) and CC-interneuron (1.0 ± 0.5 vs. 1.2 ± 0.5, P < 0.01) connections. Following ACh application, CC-CC connections and CC-interneuron connections showed an increase in the CV; at CC-interneuron connections the failure rate was also significantly increased **(Fig. 5 e; Supplementary Tab. 1)**. These changes in the EPSP properties suggest that ACh decreases the neurotransmitter release probability of intra-laminar connections established by a presynaptic L6A CC PC. Other synaptic properties, like rise time, latency and decay time, were not affected by ACh **(Supplementary Tab. 1)**.

**Fig. 5.**
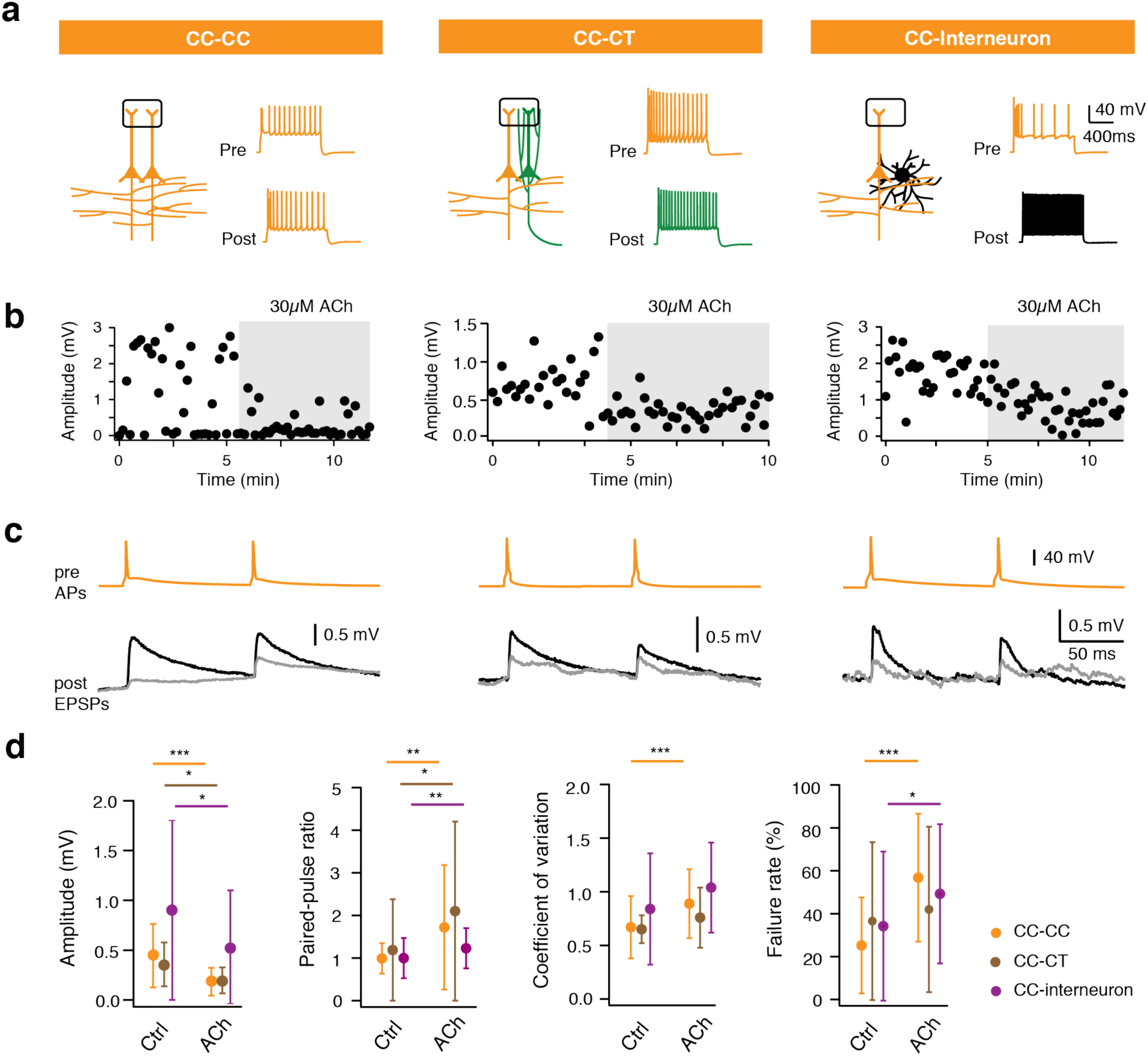
ACh-mediated reduction of presynaptic release at CC PC synapses. **(a)** Left, Schematic representation of the synaptic connections with a presynaptic CC L6A PC. CC PCs are shown in orange, the CT PC in green and the interneuron in black. Barrel structures indicate layer 4. Right, corresponding firing patterns of pre- and postsynaptic neurons of the same connection type. **(b)** Time course of EPSP amplitude changes following bath application of 30µM ACh (gray phases) in a CC-CC, a CC-CT and a CC-interneuron pair. **(c)** Overlay of average EPSPs in control (black) and ACh application (gray) phases. Presynaptic APs are shown at the top. Data are recorded from the same pairs as in (b). **(d)** The average and SD of several EPSP properties for CC-CC (n = 20), CC-CT (n = 5) and CC-interneuron (n = 9) connections are shown. *P < 0.05, **P < 0.01, ***P < 0.001 for Wilcoxon signed-rank test.

Previous studies have shown that ACh may inhibits intracortical excitatory synaptic transmission at some synaptic connections through activation of presynaptic M_4_ mAChRs^21,47,48^. To test whether the ACh-induced suppression of the efficacy of synaptic connections with a presynaptic CC L6A PC is mediated by M_4_ mAChR activation, 1 µM TRO (a selective antagonist of M_4_Rs) was co-applied with ACh (30 µM) after bath application of ACh alone. The effects of ACh on synaptic connections established by CC PCs (n = 6 pairs, comprising 2 CC-CC, 1 CC-CT and 3 CC-interneuron connections) were completely blocked by TRO. The EPSP amplitude decreased from 1.0 ± 0.8 mV to 0.3 ± 0.2 mV during ACh application and fully recovered to 1.0 ± 0.8 mV during co-application of ACh and TRO. Moreover, TRO also blocked the ACh effects on the CV (0.8 ± 0.2 for control vs. 0.9 ± 0.3 for ACh and TRO; n = 6 pairs, P = 0.75) and failure rate (27.0 ± 17.6 % for control vs. 26.3 ± 19.6 % for ACh and TRO; n = 6 pairs, P = 1.00) **(Fig. 6)**. In addition to reversing the ACh-induced increase in the PPR, TRO increased the PPR of connections established by CC PCs. Co-application of ACh and TRO resulted in a smaller PPR compared to control (1.1 ± 0.3 vs. 1.3 ± 0.4; n = 6 pairs, P < 0.05) **(Fig. 6)**. In order to isolate the presynaptic effect of ACh on L6A intra-laminar connections established by presynaptic CC PCs, a CC-CT synaptically coupled pair was recorded. The ACh-induced reduction in synaptic release probability recovered only after co-application of TRO together with PIR but not when PIR (0.5 µM) was applied alone **(Supplementary Fig. 5)**.

**Fig. 6.**
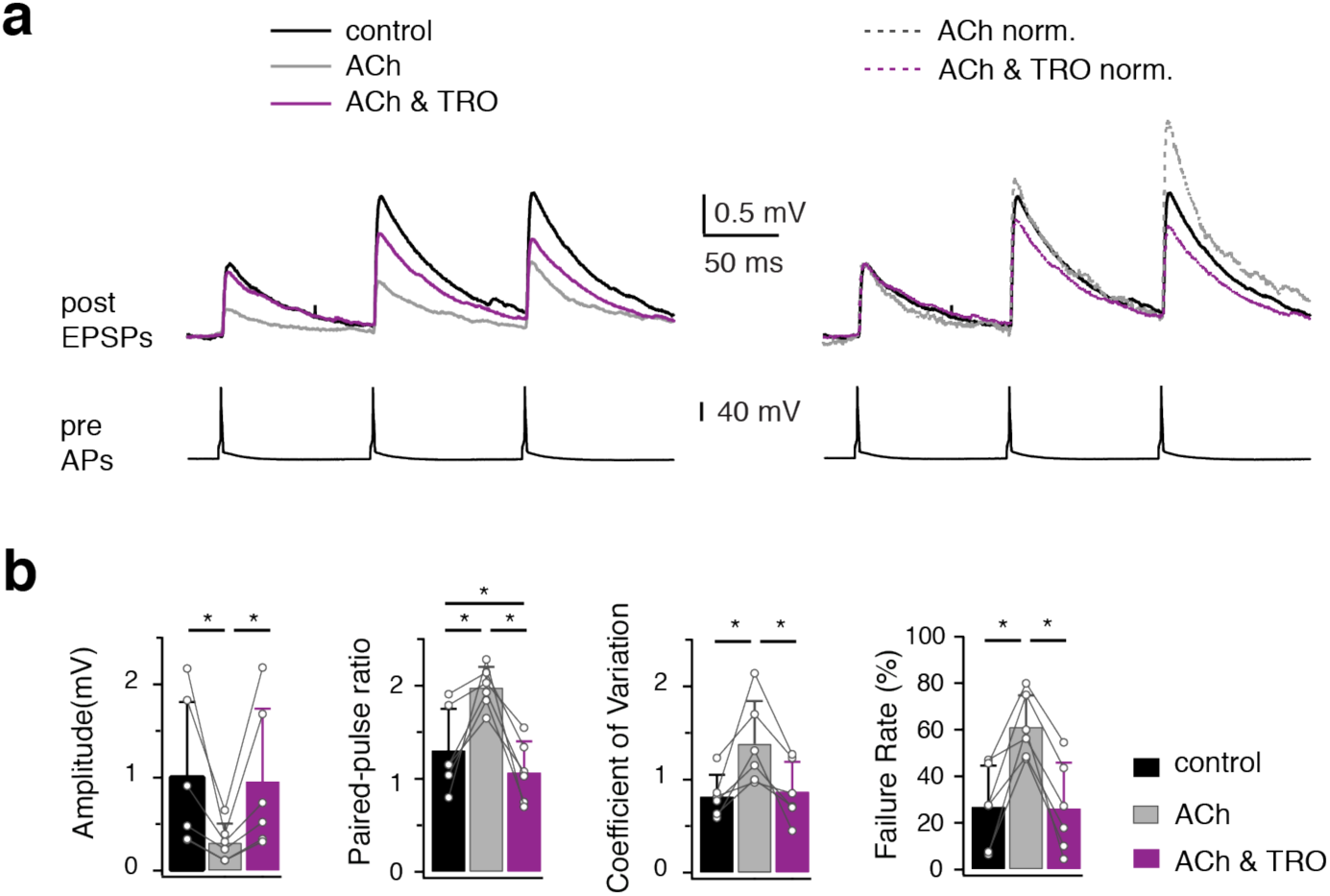
ACh decreases presynaptic release probability of CC PC via activation of M_4_ AChRs. **(a)** Left, Overlay of average EPSPs recorded in control, the presence of ACh (30µM), and of ACh & TRO (1 µM) from a representative CC-CC connection. Right, normalising the mean EPSP amplitudes obtained in ACh and ACh & TRO to the first EPSPs amplitude in control reveals changes of PPR. Presynaptic APs are shown at the bottom. **(b)** Histograms (n = 6) showing the effect of ACh and TRO blockade of ACh-induced changes on several EPSP properties including EPSP amplitude, PPR, CV and failure rate. Data were recorded from L6A synaptic connections with a presynaptic CC PC. Open circles, individual data points; bars, the average for each condition. Error bars represent SD. *P < 0.05 for Wilcoxon signed-rank test.

Because of their sparse and narrow axonal domain, L6A CT PCs rarely innervate neurons in their home layer and their intracortical synaptic connections are remarkably weak and unreliable^44,49,50^. Here we applied ACh (30 µM) to seven synaptic connections established by a presynaptic CT PC in L6A, including two CT-CT, one CT-CC and four CT-interneuron connections. In all synaptic connections established by CT PCs ACh significantly enhanced the EPSP amplitude (0.10 ± 0.08 mV vs. 0.15 ± 0.10 mV; n = 7 pairs, P < 0.05) and reduced the PPR (3.0 ± 1.7 vs. 0.8 ± 0.6; n = 7 pairs, P < 0.05) **(Fig. 7)**. The ACh-mediated reduction in the PPR suggests a presynaptic locus for synaptic modulation. Because these connections display very small unitary EPSP amplitudes and frequent failures, the signal-to-noise ratio was often too low to appropriately calculate the CV and failure rate. No significant differences were detected in rise time, decay time and latency **(Supplementary Tab. 1)**. Our findings indicate that in contrast to the inhibition of presynaptic release in L6A CC PCs, ACh enhances the synaptic efficacy of the weak connections established by a presynaptic CT PC.

**Fig. 7.**
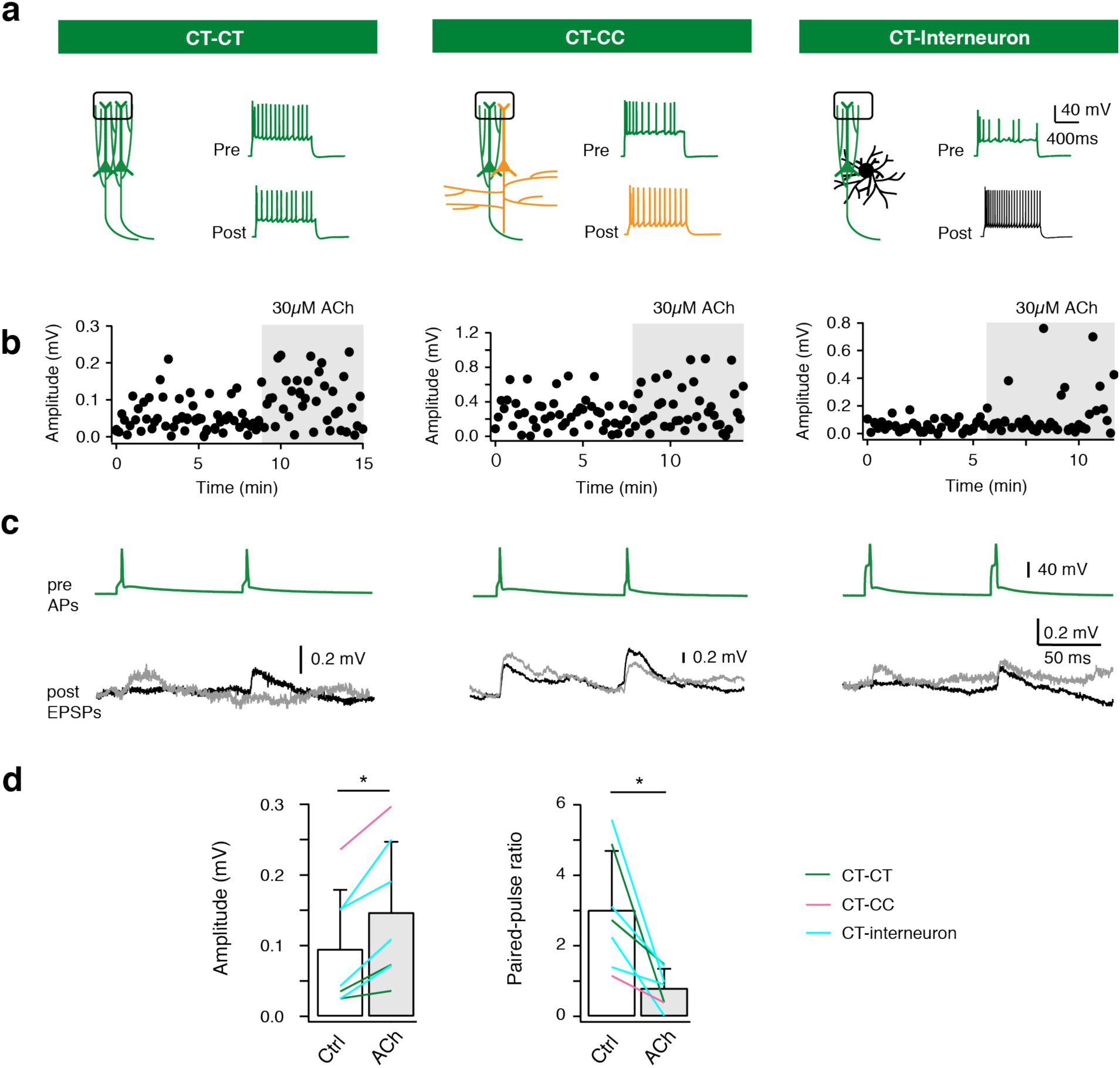
ACh enhances synaptic efficacy of L6A excitatory connections with a presynaptic CT PC. **(a)** Left, schematic representation of the synaptic connections with a presynaptic CT PC. Color code as in Fig. 5. Barrel structures indicate layer 4. Right, corresponding firing patterns of pre- and postsynaptic neurons of the same connection type. **(b)** Time course of EPSP amplitude change following bath application of 30 µM ACh in a CT-CT, CT-CC, and CT-interneuron pair. **(c)** Overlay of average EPSPs in control (black) and ACh application (gray) phases. Presynaptic APs are shown at the top. Data are recorded from the same pairs as in (b). **(d)** Summary data (n = 7) of ACh-induced changes in first uEPSP amplitude and paired-pulse ratio for L6A excitatory pairs with a presynaptic CT L6A PC. Bars indicate the average for each condition; Error bars represent SD. *P < 0.05 for Wilcoxon signed-rank test.

To determine the AChR subtype that mediates the increase in synaptic efficacy at these connections, we tested whether the selective antagonists of the M_1_Rs (PIR), homomeric the α_7_ subunit-containing nAChRs (methyllycaconitine, MLA) and the heteromeric α_4_ß_2_ subunit-containing nAChRs (DHßE) could block the effect of ACh on synaptic connections with a presynaptic CT PC. While PIR and MLA had no effect, DHßE blocked the increase of EPSP amplitude and decrease of PPR **(Fig. 8)**. This indicates that the ACh-induced enhancement of synaptic efficacy is induced by activation of α_4_ß_2_ subunit-containing nAChRs in presynaptic CT PCs.

**Fig. 8.**
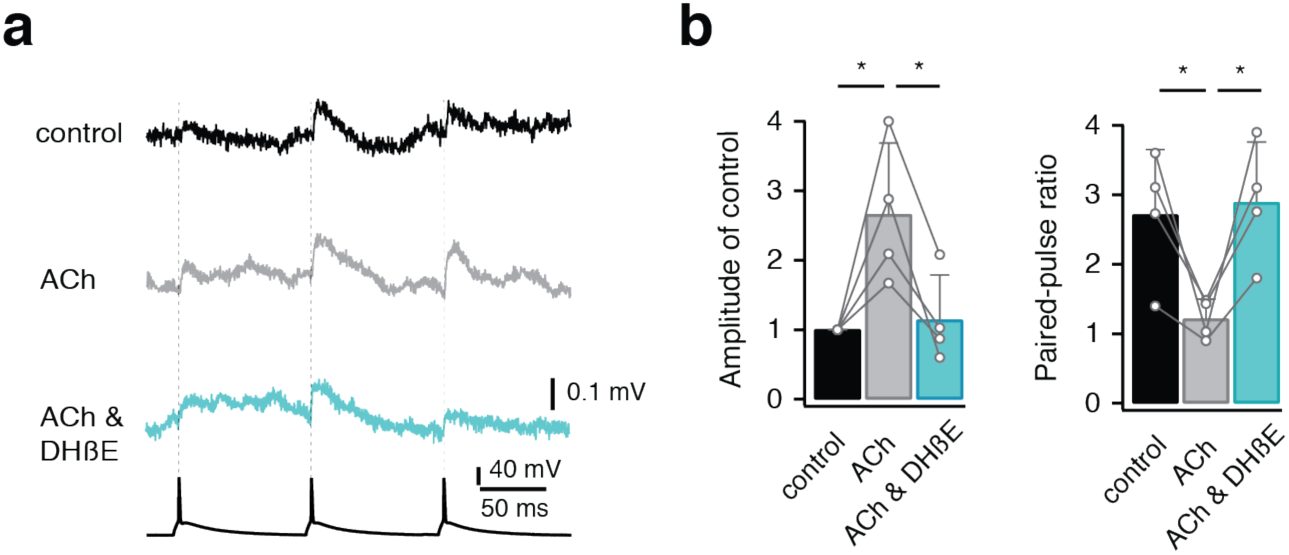
ACh increases presynaptic release probability of CT PC via activation of α_4_β_2_ nAChRs. **(a)** Average EPSPs recorded in control, the presence of ACh (30µM), and of ACh & DHßE (10 µM) from a representative CT-CT connection. Gray phase, bath application of 30µM ACh; Turquoise phase, co-application of ACh and DHßE. The presynaptic APs are shown at the bottom. **(b)** Histograms (n = 4) showing the effect of ACh and DHßE blockade of ACh-induced changes in first uEPSP amplitude and paired-pulse ratio. Data were recorded from L6A synaptic connections with a presynaptic CT PC. Open circles, individual data points; bars, the average for each condition. Error bars represent SD. *P < 0.05 for paired Student’s t-test.

## Discussion

We investigated the cholinergic modulation of CT and CC PCs in layer 6A of the barrel cortex. We showed that (i) low concentrations of ACh differentially modulate the L6A microcircuitry by persistently depolarizing CT but hyperpolarizing CC L6A PCs. These effects are monophasic and mediated via M_1_ and M_4_ mAChRs, respectively; (ii) a nicotinic ACh response was observed exclusively in CT PCs only when a high ACh concentration was applied. In addition, (iii) low concentrations of ACh increases the frequency of miniature EPSCs via presynaptic nAChRs in L6A CT but decreases that of CC PCs via M_4_Rs. To better understand the effects of ACh on intralaminar synaptic transmission, recordings were performed from synaptically coupled L6A PC pairs. We found that (iv) in neuronal connections with a presynaptic CC PC the neurotransmitter release probability was reduced via activation of M_4_Rs but (v) increased in connections with a presynaptic CT L6A neuron by α_4_β_2_ nAChR activation. Our results reveal that two functionally and morphologically distinct subpopulations of L6A PCs are affected differentially by ACh acting on both mAChRs and nAChRs.

### Synergistic modulation of L6A PCs by mAChRs and nAChRs

In a number of studies investigating the nAChR response of L6 PCs in different cortical areas only high ACh concentration (≥ 1 mM) have been applied because the ACh affinity of nAChRs is substantially lower than that of mAChRs^23,36,37,39^. Under this condition any mAChR effect is almost entirely masked by the strong nicotinic response so that any involvement of mAChRs has been explicitly ruled out. Here we demonstrate for the first time that mAChRs play crucial roles in both pre- and postsynaptic modulation of L6A PC activity. Both the pre- and postsynaptic effects of mAChRs are already present at low ACh concentrations (1-10 µM) suggesting that neuromodulation via mAChRs is tonically present and mediated by volume transmission^51,52^. On the other hand, only a high concentration of ACh (EC_50_ of ∼1 mM) could effectively depolarize CT L6A PCs via nAChRs, but an up-regulation of presynaptic vesicle release via α_4_β_2_ subunit-containing nAChRs was already observed in the presence of 30 µM ACh. This α_4_β_2_ nAChR-mediated effect was found for spontaneous excitatory synaptic activity as well as for CT-formed monosynaptic connections, suggesting that presynaptic nAChR are expressed on synaptic boutons of both CT L6A axons and other glutamatergic afferents (e.g. from thalamus or other cortical regions). Single channel currents from nAChRs can be activated already by low concentrations of ACh^53,54^. Because of the electrotonic compact structure of a presynaptic bouton even the opening of a few nAChR receptor channels may result in a depolarisation that is sufficiently strong to increase the open probability of presynaptic Ca^2+^ channels and hence the neurotransmitter release.

Furthermore, in some L6 PCs the α_5_ nAChR subunit co-assembles with the α_4_ and ß_2_ subunits^23,36^. Nicotinic AChRs containing the α_4_, ß_2_ and α_5_ subunits have a higher Ca^2+^ permeability than those composed of α_4_ and ß_2_ nAChR subunits alone^55^. In the presynaptic terminals, Ca^2+^ entry via α_4_ß_2_α_5_ nAChRs into the presynaptic bouton could enhance neurotransmitter release provided these channels are located sufficiently close to the release site.

### Cholinergic activation by endogenous ACh in neocortical L6

In the cortex, ACh levels change dramatically during different stages of waking and sleep^8,9^. It has been suggested that high ACh levels serve to enhance the response to sensory stimuli by increasing the strength of afferent input while low concentration of ACh contributes in the consolidation of encoded information^56^. Cholinergic signalling has been described to occur via a volume release mechanism^52^, which is slow and unspecific. Volume release of ACh reaches concentrations in a low micromolar range, spreads widely over neocortical layers and activates predominantly mAChRs. In addition, cholinergic synapses have been identified particularly in deep layers of neocortex and less so in superficial cortical layers^20,36,57^. At these cholinergic synapses, ACh reaches a high concentration in the synaptic cleft, thereby activating postsynaptic nAChRs in L6 PCs^36^. Because cholinergic synapses in the neocortex are small^58^, ACh released into the synaptic cleft may spill over into the perisynaptic space. Subsequently, the extra-synaptic AChRs on presynaptic boutons of CT PCs are activated resulting in an increase of release probability (**Fig. 7**). Thus, nAChRs and mAChRs act on different time scales and at different neurotransmitter concentrations, resulting in a striking complexity of the cholinergic modulation of neocortical signaling.

### The cell type-specific effect of ACh in L6A

ACh has been shown to induce a persistent depolarization of L2/3 and L5 PCs but a hyperpolarization of excitatory L4 neurons^21,43,59^. Here, we demonstrate that ACh modulates PCs not only in a layer-specific but also a cell type-specific way that can be attributed to a cell type-dependent expression of mAChRs (**Fig. 9**). In L6A of barrel cortex, ACh hyperpolarizes CC PCs but depolarizes CT PCs via activation of M_4_Rs and M_1_Rs, respectively. The action potential firing frequency was decreased by ACh in CC PCs but increased in CT PCs, thereby modulating the excitability and signal propagation in L6A PCs in a cell-specific manner. In addition, CT L6A PCs but not CC PCs showed a strong α_4_β_2_ nAChR-mediated response (**Fig. 9**). This is consistent with previous findings in L6 of prefrontal cortex that regular spiking neurons have a larger nicotinic receptor-mediated inward current following ACh application when compared with bursting neurons^37^. A cell type-specific neuromodulation was also discovered previously in deep layers of medial prefrontal cortex for neuromodulators such as noradrenaline, dopamine and adenosine^60-64^.

**Fig. 9.**
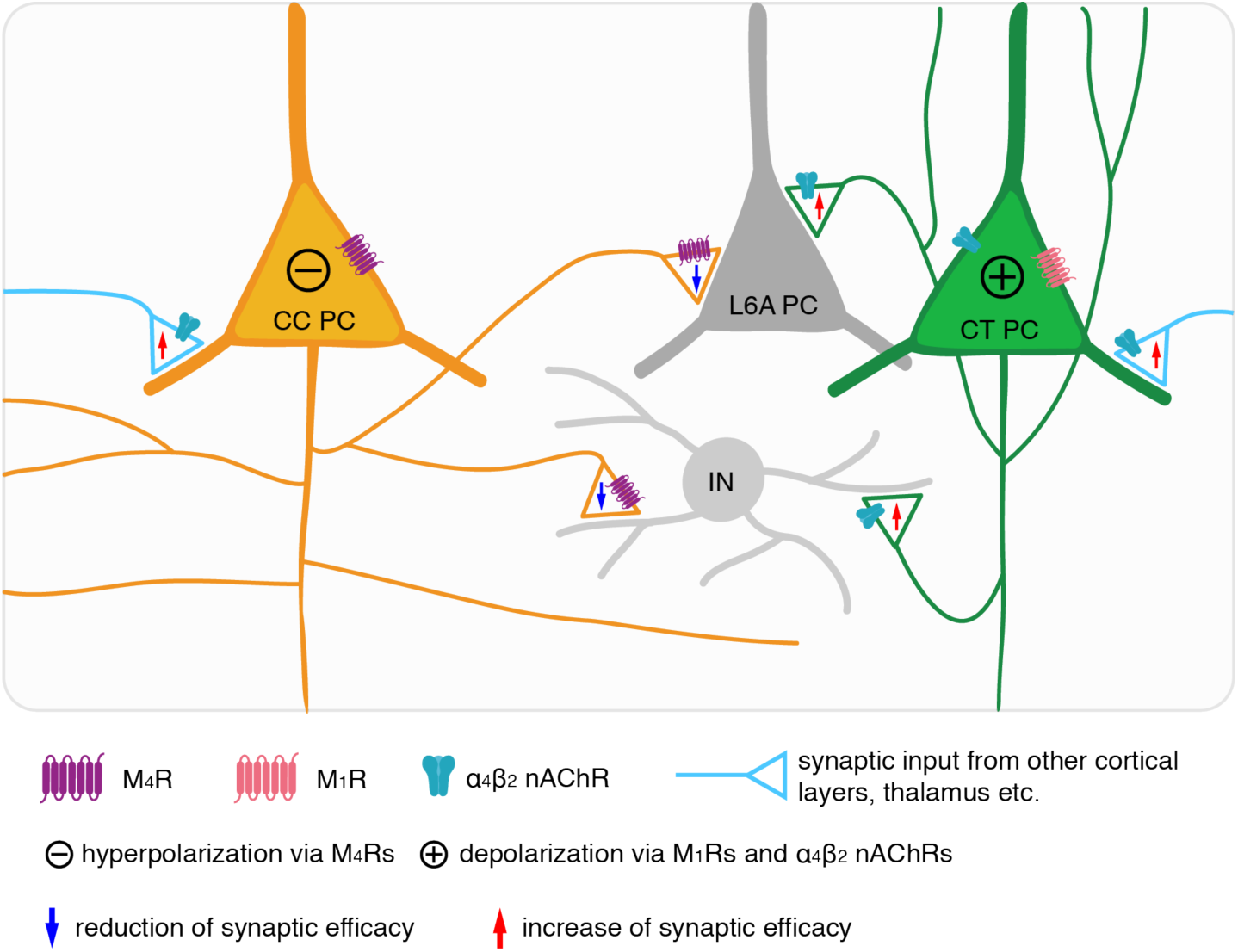
Cholinergic actions on muscarinic and nicotinic receptors in L6A PCs. Schematic summary shows cholinergic modulation of L6A microcircuits in rat barrel cortex. ACh affects membrane excitability and presynaptic release probability of L6A PCs via activating muscarinic and/or nicotinic AChRs. Because of the cell type-specific distribution of AChRs at pre- and postsynaptic sites, CC and CT L6A PCs are affected differentially by ACh. The presynaptic CC and CT L6A PC are shown in orange and green, respectively. The postsynaptic L6A PC and interneuron are shown in gray.

By studying miniature spontaneous activity of L6A CC and CT PCs, we found that ACh both increases the excitatory synaptic release onto CC and CT PCs by activation of α_4_β_2_ nAChRs (**Fig. 9**). Because CT PCs express α_4_β_2_ nAChRs, an increase in spontaneous activity may result from the enhanced release probability at CT L6 PC boutons; however, the intracortical axon density of these PCs is low so that their contribution to the spontaneous mEPSC frequency is minimal. On the other hand, activation of nAChRs increases thalamocortical transmission onto L3, L4 and L5 neocortical neurons^47,65,66^. Thus, the increased excitatory transmission onto L6A PCs is probably resulting to a large degree from a higher release probability at thalamocortical and less so from intracortical synapses. In addition, CC PCs receive more intra-laminar inputs than CT PCs, which can be suppressed by ACh via M_4_Rs. Therefore, the ACh-induced reduction of mEPSC frequency in CC PCs could be a combinatorial effect on thalamocortical and intracortical transmission.

It has been proposed that ACh increases the signal-to-noise ratio (SNR) of sensory signaling by selectively enhancing thalamocortical inputs over intracortical synaptic transmission^47,67,68^. ACh has been found to suppress the efficacy of excitatory intracortical connections in different layers including L2/3, L4 and L5^21,47,48^. Here, a differential cholinergic modulation of presynaptic neurotransmitter release was observed in CC and CT L6A PC types. ACh suppresses synaptic transmission in excitatory L6A connections with presynaptic CC PCs through activation of M_4_Rs but potentiates connections with a presynaptic CT PCs via presynaptic α4ß2 nAChRs (**Fig. 9**); no α7 nAChR or M_1_R effect on synaptic transmission was observed.

In hippocampus and some subcortical structures such as the ventral tegmental area, glutamatergic synapses are known to be facilitated by nAChRs located on presynaptic terminals^69,70^. However, very few studies demonstrate an ACh-mediated enhancement of intracortical excitatory synaptic transmission. Recently, it has been shown that excitatory synaptic transmission between PCs and somatostatin-positive interneurons in layer 2 of mouse barrel cortex is increased by ACh via activating nAChRs^71^. Although an increase of synaptic efficacy was observed in connections with a presynaptic CT PCs, this type of synaptic connections are rare, generally weak and very unreliable^49,50^. Therefore we propose that ACh mainly acts on CT PCs not primarily by increasing intracortical synaptic transmission but rather by facilitating corticothalamocortical feedback; this facilitation will occur already at ACh levels in the low micromolar range.

## Acknowledgement

We thank Werner Hucko for excellent technical assistance and Dr. Dr. Karlijn van Aerde for custom-written macros in Igor Pro software. We warmly thank Dr. Chao Ding for helpful discussions and Dr. Vishalini Emmenegger for proofreading the manuscript.

## Author contributions

D.F., G.R. and D.Y. designed the research; D.Y., R.G. and G.Q. performed experiments and data analysis; D.Y., D.F. and G.R. wrote the manuscript. The authors declare no conflicts of interest.

## Materials and Methods

### Slice preparation and solutions for electrophysiology

All experiments involving animals were performed in accordance with the EU Directive 2010/63/EU, the German animal welfare act and the guidelines of the Federation of European Laboratory Animal Science Association (FELASA). Wistar rats (Charles River) were maintained on a 12/12-hr light-dark cycle from 7 AM to 7 PM. Rats aged 17-21 postnatal days (P17-21, both sexes) were lightly anaesthetized with a concentration < 0.1% of isoflurane and then decapitated. The brain was quickly removed and transferred into ice-cold artificial cerebrospinal fluid (ACSF) containing a high Mg^2+^- and a low Ca^2+^-concentration (4 mM MgCl_2_ and 1 mM CaCl_2_) to reduce synaptic activity and bubbled continuously with carbogen (95% O_2_ and 5% CO_2_). It was then placed on the ramp of a slope of 10° and were cut at an angle of 50° to the midline^72^. Thalamocortical slices were cut at 350 µm thickness using a high vibration frequency and incubated for 30-60 minutes at room temperature (21-24°C) in slicing solution. During whole-cell patch clamp recordings, slices were continuously perfused with a perfusion solution containing (in mM): 125 NaCl, 2.5 KCl, 25 D-glucose, 25 NaHCO_3_, 1.25 NaH_2_PO_4_, 2 CaCl_2_, 1 MgCl_2_, 3 myo-inositol, 2 sodium pyruvate and 0.4 ascorbic acid, bubbled with carbogen and maintained at a temperature of 30-33°C. Patch pipettes were filled with an internal solution containing (in mM): 135 K gluconate, 4 KCl, 10 HEPES, 10 phosphocreatine, 4 Mg-ATP and 0.3 GTP (pH 7.4, 290-300 mOsm). To stain the patched neurons, biocytin was added at a concentration between 3-5 mg/ml to the pipette solution; a recording time of ∼15-30 minutes was necessary for biocytin to diffuse into the dendrites and axons of the recorded cells^73,74^. No biocytin was added to the internal solution of ‘searching’ pipettes used during searching for synaptic connections.

### Cell identification

Slices were placed in the recording chamber under an upright microscope (fitted with 4x plan/ 0.13 numerical aperture and 40x water immersion/0.80 NA objectives; Olympus, Tokyo, Japan) with the pial surface pointing forward. The cortical layers and the barrel field were visualized at 4x magnification; the barrels can be identified in L4 as narrow dark stripes with evenly spaced, light ‘hollows’ and were visible in 6-8 consecutive slices. L6A neurons were identified in the upper 60% of layer 6 at 40x magnification using infrared differential interference contrast (IR-DIC) microscope^75,76^. Putative PCs and interneurons were differentiated on the basis of their intrinsic action potential firing pattern during recording and after histological processing by their morphological appearance.

### Electrophysiological recordings

Whole-cell patch clamp recordings from L6A neurons were performed at 30-33°C for an optimal oxygenation. Patch pipettes were pulled from thick-wall borosilicate capillaries (outer diameter: 2 mm; inner diameter: 1 mm) to a final resistance of 6-10 MΩ. Recordings were made using an EPC10 amplifier (HEKA, Lambrecht, Germany), sampled at 10 kHz, and filtered at 2.9 kHz using the Patch-master software (HEKA). Neurons were selected randomly and excluded from the analysis when their whole-cell series resistance exceeded 40 MΩ (50 MΩ for neurons from paired-recodings) or their resting membrane potential was more depolarized than -50 mV immediately after rupturing the cell membrane. The resting membrane potential of L6A excitatory neurons was continuously recorded in the current clamp mode to monitor changes in amplitude.

Miniature spontaneous events were recorded in voltage-clamp mode and changes in mEPSC frequency and amplitude were analyzed. Recordings of L6A excitatory neurons were made in the presence of tetrodotoxin (TTX, 0.5 µM) and gabazine (10 µM) to inhibit AP firing and inhibitory postsynaptic currents (IPSCs), respectively. During recordings, the holding potential was set at - 70mV.

Because the connectivity of L6A neurons was low compared to other intra-laminar connections in rat barrel cortex, we followed the ‘searching procedure’ described previously after patching a putative postsynaptic neuron^74,77^. A monosynaptic connection can be found by patching multiple cells in ‘loose cell-attached’ mode. When the AP resulted in a unitary excitatory postsynaptic potential (uEPSP) in the postsynaptic L6A neuron, this presynaptic neuron was repatched with a new pipette filled with biocytin containing internal solution. APs were elicited by current injection in the presynaptic neurons and the postsynaptic response were recorded in whole cell (current clamp) mode, the effects of ACh on unitary EPSPs were then tested.

### Drug Application

ACh (1 µM-10 mM) was bath applied via the perfusion system or puff applied through a patch pipette (tip diameter: 1-2 µm) connected to a PDES-02D device (npi electronic GmbH, Tamm, Germany). The puff pipette was placed at 10-20 µm from the same recorded neuron and a brief low pressure was applied for about 1 s. Mecamylamine (10 µM), atropine (200 nM-20 µM), pirenzepine (0.5 µM), tropicamide (1µM), dihydro-ß-erythroidine (DHßE) (10 µM), methyllycaconitine (MLA) (10µM), tetrodotoxin (TTX) (0.5 µM) and gabazine (10 µM) were all bath applied; drugs were purchased from Sigma-Aldrich (Steinheim, Germany) or Tocris (Bristol, UK).

### Histological staining

After single cell or paired recordings, brain slices containing biocytin-filled neurons were processed as described previously^73^. Slices were fixed at 4°C for at least 12 hours in 100 mM phosphate buffer (PB, PH 7.4) solution containing 4% paraformaldehyde (PFA) and then incubated in 0.1% triton X-100 solution containing avidin-biotinylated horseradish peroxidase (Vector ABC staining kit, Vector Lab. Inc., Burlingame, USA). The reaction was catalyzed using 3,3′-diaminobenzidine (Sigma-Aldrich, St. Louis, MO, USA) as a chromogen. Slices were again rinsed with 100 mM PB solution several times, followed by slow dehydration using ethanol and xylene. After embedding in Eukitt medium (Otto Kindler GmbH, Freiburg, Germany), the dendritic and axonal structures were clearly visible.

Immunofluorescence staining was performed for the identification of molecular markers expressed in L6A PCs. During electrophysiological recordings, Alexa Fluor^®^ 594 dye (1:500, Invitrogen, Darmstadt, Germany) was added to the internal solution for post hoc identification of patched neurons. After recording, slices (350 µm) were fixed with 4% PFA in 100mM PBS for at least 24 hours at 4°C and then permeabilized in 1% milk power solution containing 0.5% Triton X-100 and 100 mM PBS. Primary and secondary antibodies were diluted in the permeabilization solution (0.5% Triton X-100 and 100 mM PBS) shortly before experiments. For single cell-FoxP2 staining, slices were incubated overnight with Goat-anti-FoxP2 primary antibody (1:500, Santa Cruz Biotechnology, Heidelberg, Germany) at 4°C and then rinsed thoroughly with 100 mM PBS. Subsequently, slices were treated with Alexa Fluor^®^ secondary antibodies (1:500) for 2-3 hours at room temperature in the dark. After being rinsed in 100 mM PBS the slices were embedded in Moviol. The fluorescence images were taken using the Olympus CellSens platform. The position of the patched neurons were identified by the conjugated Alexa dye, so that the expression of FoxP2 could be tested in biocytin-stained neurons. After acquiring fluorescent images, slices were incubated in 100 mM PBS overnight and were processed for subsequent morphological analysis. Co-immunostaining of FoxP2 and M_4_Rs (Rabbit-anti-M_4_Rs, 1:500, Abbexa, Cambridge, UK) were performed with 150 µm thin brain slices following the procedure described above.

### Morphological reconstructions

3D reconstructions of L6A excitatory and inhibitory neurons or synaptically coupled neuron pairs labelled with biocytin were made using the NEUROLUCIDA^**®**^ software (MicroBrightField Inc., Williston, VT, USA) and Olympus BX61 microscopy at 1000 X magnification. Slices were selected to be reconstructed only if the labeling quality was high and the background staining was low. Barrel borders, demarcation of different layers, pial surface and white matter were delineated during reconstructions. The cell body, the axonal and dendritic branches were reconstructed manually under constant visual inspection to detect even small collaterals. Corrections for shrinkage were performed in all spatial dimensions (factor 1.1 in the x and y axes, factor 2.1 in the z axes) 73. Analysis of 3D reconstructed neurons was done with NEUROEXPLORER^**®**^ software (MicroBrightField Inc., Williston, VT, USA).

The neuronal polarity of reconstruction was calculated with NEUROEXPLORER^®^ software using cubic spline smoothing. The dendritic and axonal length was averaged for each of the 120 “3° sectors” around the soma. Data were recalculated, plotted in angular subdivision around the soma and polar plots were made with Grapher software (GoldenSoftware, Colorado, USA). The radian depicts degree in angles (°) with 0° towards the pial surface, 90° towards the posterior-median axis, 180° towards the white matter, and 270° towards the anterior-lateral axis.

### Data analysis

Custom written macros for Igor Pro 6 (WaveMetrics, Lake Oswego, USA) were used to analyze the recorded electrophysiological signals. The miniature spontaneous activity was analyzed using the program SPCN (http://www.spacan.net). A threshold of 5 pA was set manually for detecting mEPSC events, which is at least 2.5 fold larger than the noise level (< 2pA). No noise filtration was applied before data analysis.

The synaptic properties were evaluated as described in the previous studies^77^. First, all sweeps were aligned to their corresponding presynaptic AP peaks and an average sweep was generated as the mean uEPSP. The EPSP amplitude was calculated as the difference between the mean baseline amplitude and maximum voltage of the postsynaptic event. The paired pulse ratio was defined as the second uPSP amplitude divided by the first uPSP amplitude of the mean uPSP elicited by paired APs with a stimulation frequency of 10 Hz. Failures were defined as events with amplitudes <1.5× the standard deviation (SD) of the noise within the baseline window and the failure rate refers to the percentage of failures. The coefficient of variation (CV) was calculated as the SD divided by the mean uEPSP amplitude.

### Statistical tests

For all data, the mean ± SD was given. To assess the differences between two paired groups under different pharmacological conditions, Wilcoxon signed rank test was performed. The paired Student’s *t* test was used when n = 4 for the paired samples and the Mann-Whitney *U*-test was used when the sample size was different between two groups. Statistical significance was set at P < 0.05, n indicates the number of neurons or pairs analyzed.

