## Supplementary materials for "Cell Type-Specific Modulation of Layer 6A Excitatory Microcircuits by Acetylcholine in Rat Barrel Cortex"

*Corresponding author: Dirk Feldmeyer

### Supplementary Materials

Containing 1 table and 5 figures with legends

| | CC-CC (n = 20 pairs) | CC-CT (n = 5 pairs) | CC-Interneuron (n = 9 pairs) | CT-connections (n = 7 pairs) |
|---|---|---|---|---|
| | **Control** | | | |
| Amplitude (mV) | 0.45 ± 0.32 | 0.35 ± 0.22 | 0.90 ± 0.90 | 0.10 ± 0.08 |
| PPR | 1.0 ± 0.4 | 1.2 ± 0.7 | 1.0 ± 0.5 | 3.0 ± 1.7 |
| CV | 0.7 ± 0.3 | 0.7 ± 0.1 | 0.8 ± 0.5 | - |
| Failure rate (%) | 25.2 ± 22.3 | 36.5 ± 36.8 | 34.3 ± 34.8 | - |
| Rise time (ms) | 1.5 ± 0.5 | 1.8 ± 1.8 | 1.0 ± 0.5 | 1.1 ± 0.4 |
| Latency (ms) | 1.8 ± 0.9 | 1.7 ± 0.5 | 1.1 ± 0.5 | 2.3 ± 0.6 |
| Decay time (ms) | 37.4 ± 17.5 | 33.7 ± 15.2 | 23.2 ± 15.2 | 20.4 ± 19.0 |
| | **ACh (30µM)** | | | |
| Amplitude (mV) | *** 0.19 ± 0.14 | * 0.19 ± 0.13 | * 0.52 ± 0.58 | * 0.15 ± 0.10 |
| PPR | ** 1.5 ± 0.7 | * 2.1 ± 1.6 | ** 1.2 ± 0.5 | * 0.8 ± 0.6 |
| CV | *** 0.9 ± 0.3 | 0.8 ± 0.3 | 1.0 ± 0.4 | - |
| Failure rate (%) | *** 56.8 ± 29.8 | 42.0 ± 38.6 | * 49.3 ± 32.5 | - |
| Rise time (ms) | ** 1.2 ± 0.6 | 1.6 ±1.6 | 1.0 ± 0.4 | 1.6 ± 0.7 |
| Latency (ms) | 1.8 ± 1.0 | 1.6 ± 0.5 | 1.3 ± 0.5 | 2.2 ± 1.6 |
| Decay time (ms) | 31.1 ± 16.0 | 36.0 ± 24.2 | 16.1 ± 10.6 | 30.9 ± 28.0 |

**Supplementary Tab. 1 uEPSP properties of L6A synaptic connections under control and 30 µM acetylcholine conditions.**

Italic bold font indicates significant differences to control; *P < 0.05, **P < 0.01, ***P < 0.001 for Wilcoxon signed-rank test. CT- connections (n = 7) comprise 2 CT-CT, 1 CT-CC and 4 CT-interneuron connections.

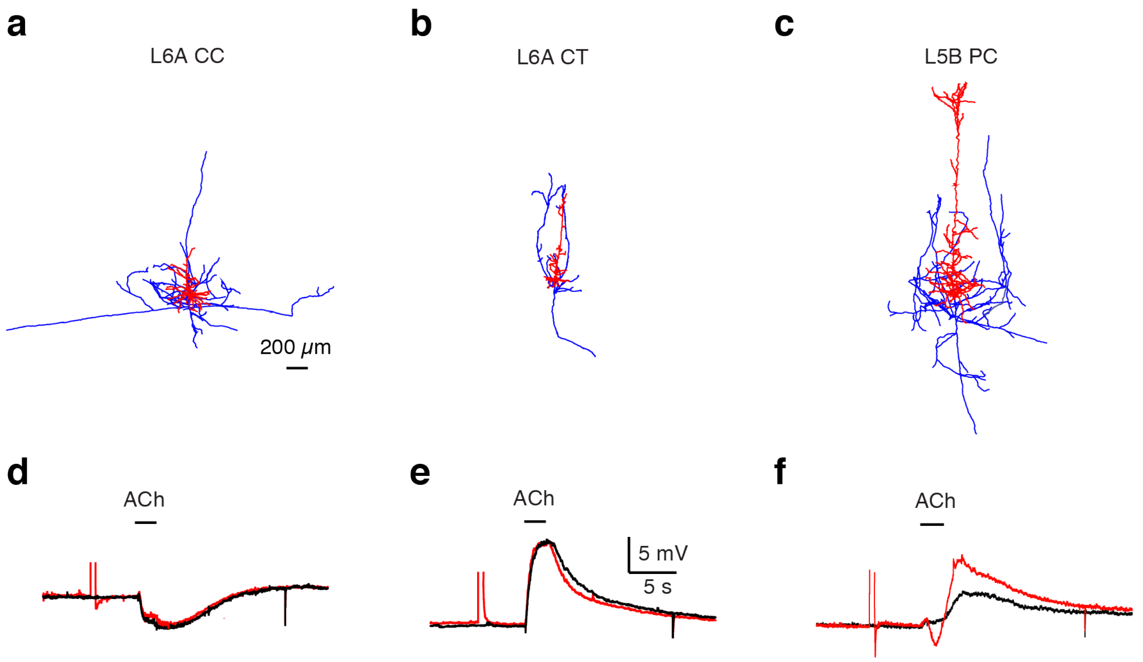


**Supplementary Fig.1 ACh-mediated hyperpolarization and depolarization are monophasic.**

**(a-c)** Morphological reconstructions of representative CC-like **(a)**, CT-like L6A pyramidal cell **(b)** and L5 pyramidal cell **(c)**.

**(d-f)** Current clamp recordings with (red) and without (black) suprathreshold depolarizing current pulse before ACh application (30µM) are recorded from the same neuron in a, b and c, respectively.

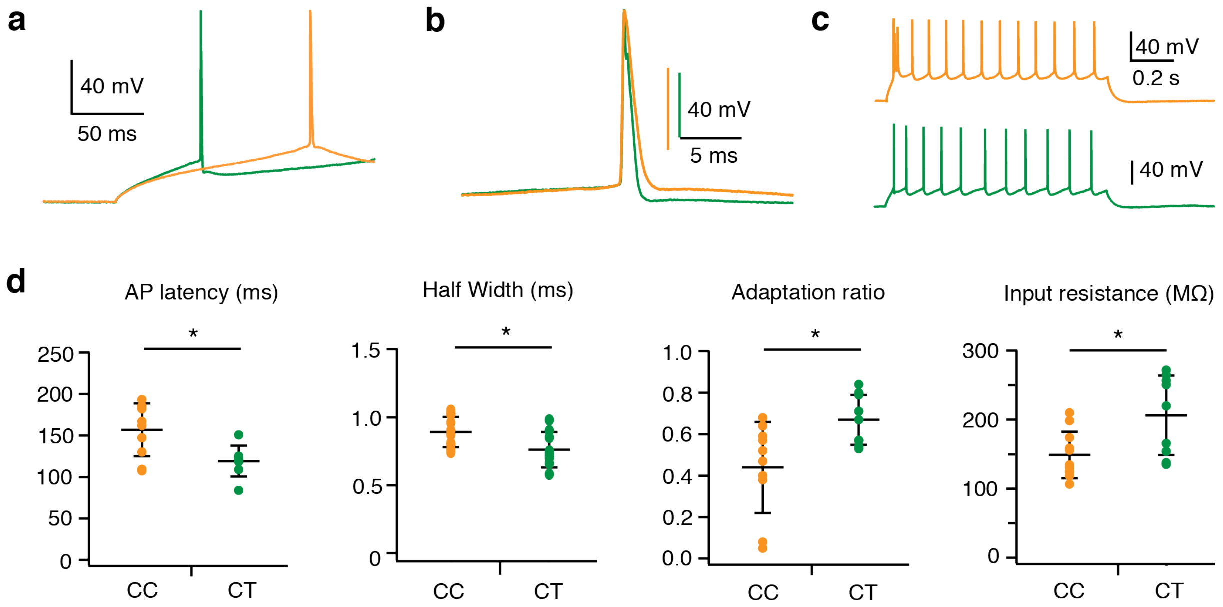


**Supplementary Fig. 2 Electrophysiological differences between CC and CT-like pyramidal cells.**

**(a)** Overlay of action potentials (APs) evoked by rheobase current from representative CC (orange) and CT (green) pyramidal cells illustrating the difference in AP time.

**(b)** Higher magnifications of the APs in **(a)** displays the difference in AP halfwidth.

**(c)** The firing patterns of the same neurons in **(a)**. An initial burst was detected in the firing pattern of CC-like pyramidal cell.

**(d)** Histograms comparing the AP latency, AP half width, Adaptation ratio (2nd ISI /10th ISI) and input resistance for CC and CT like pyramidal cells. n = 11 for CC pyramidal cells and n = 9 for CT pyramidal cells. *P < 0.05 for the Mann Whitney u-test.

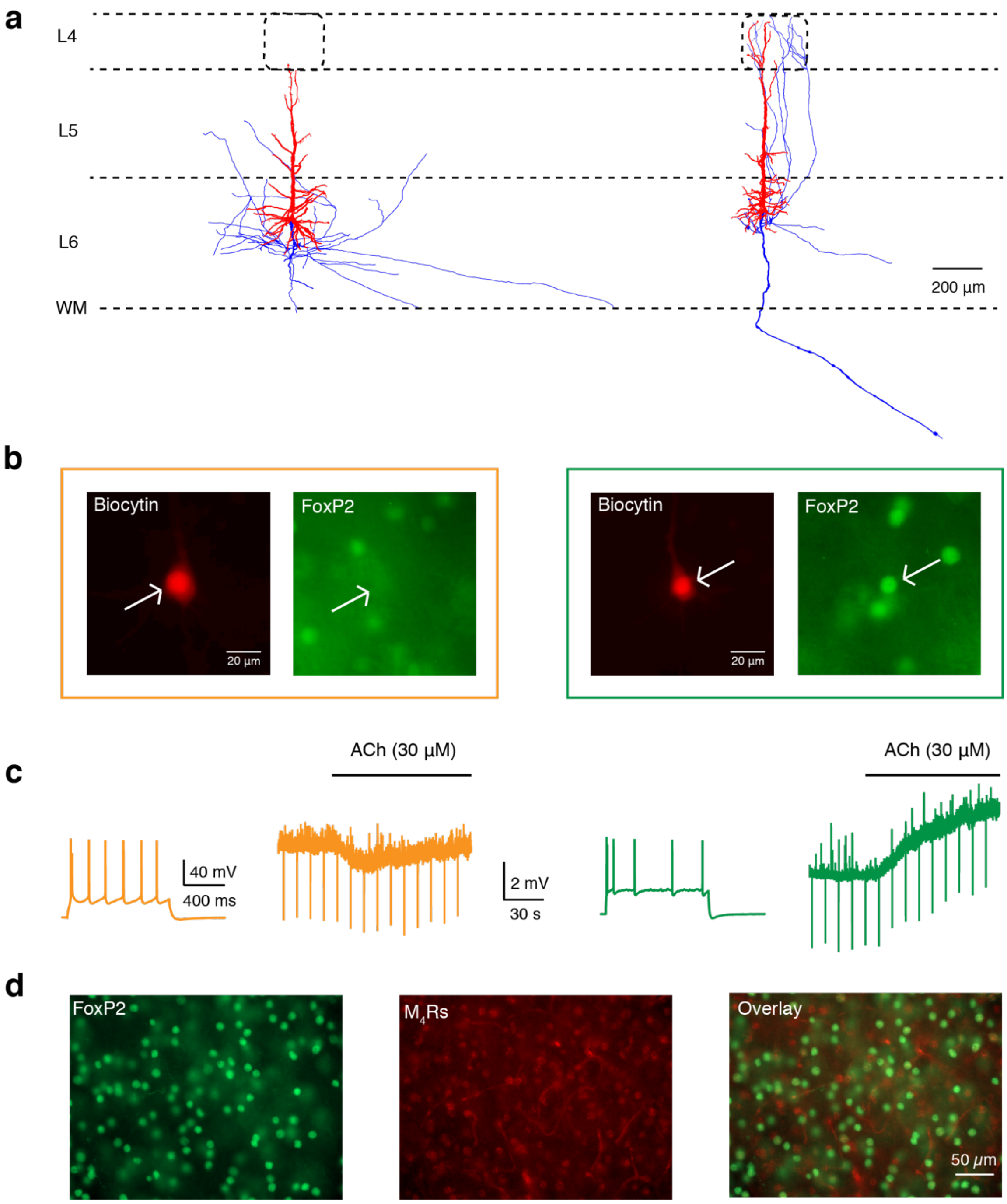


**Supplementary Fig. 3 Molecular marker expression of L6A pyramidal cells in rat barrel cortex.**

**(a)** Morphological reconstructions of a CC-like (left) and a CT-like (right) pyramidal cell. Axon is labeled in blue, soma and dendrites in red.

**(b)** The same neurons from **(a)** are recorded using whole-cell patch-clamp technique with filling biocytin coupled to Alexa 594 (red) to identify the neuronal location and morphology. The co-expression of FoxP2 (green) is tested.

**(c)** Corresponding firing patterns and cholinergic responses (30µM) of CC (left) and CT-like pyramidal cell (right) from **(a)** are shown.

**(d)** Comparison of FoxP2 immunostaining (left, green) and M4 muscarinic receptors staining (middle, red) on the same sections of cortical layer 6A in rat barrel cortex. Superimposed image is shown on the right.

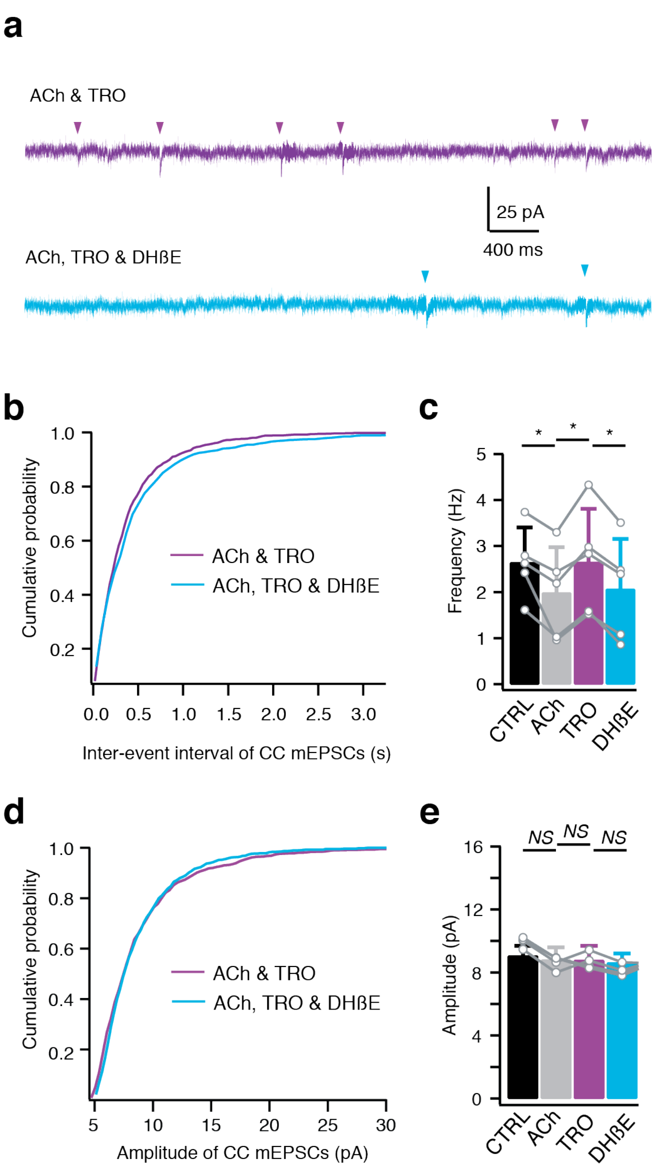


**Supplementary Fig. 4 ACh increases miniature spontaneous activity in CC PCs via α4β2 nAChRs.**

**(a)** Representative voltage-clamp recordings of a CC-like pyramidal cell following co-bath application of ACh (30µM) and different receptor antagonists. Applying of 10µM DHßE decreased frequency of mEPSC events. Traces of inward mEPSCs were recorded in the presence of TTX (0.5 µM) and GABAzine (10 µM) with a holding potential of -70 mV.

**(b)** Cumulative distributions of mEPSC inter-event interval recorded in L6A CC cells (n = 5) following co-bath application of ACh and different receptor antagonists.

**(c)** Histograms of mEPSCs frequency recorded in L6A CC cells under control, ACh, ACh & TRO and ACh & TRO & DHßE conditions. n = 5, * $P < 0.05$ for Wilcoxon signed-rank test. Error bars represent SD.

**(d)** Cumulative distributions of mEPSC amplitude recorded in L6A CC cells (n = 5) following co-bath application of ACh and different receptor antagonists.

**(e)** Histograms of mEPSCs amplitude recorded in L6A CC cells under control, ACh, ACh & TRO and ACh & TRO & DHßE conditions. n = 5, not significant (NS) for Wilcoxon signed-rank test. Error bars represent SD.

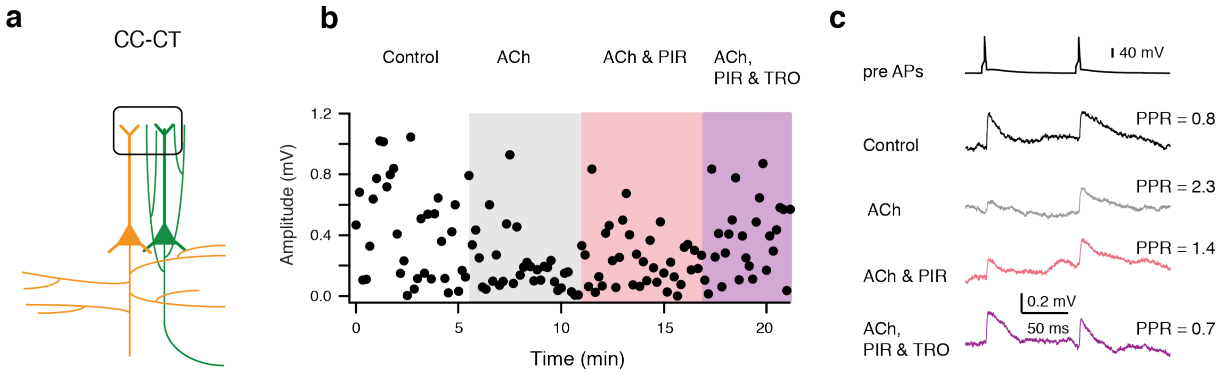


**Supplementary Fig. 5 Pharmacological experiments on a CC-CT synaptically coupled L6A pair.**

**(a)** Schematic diagram of a CC-CT connection. The presynaptic CC PC is shown in orange and the postsynaptic CT PC in green. Barrel structure indicates layer 4.

**(b)** Time course of first EPSP amplitude changes. Gray phase, application of ACh (30μM); Pink phase, co-application of ACh and PIR (0.5μM); Purple phase, co-application of ACh, PIR and TRO (1μM). Data are recorded from the pair shown in (a).

**(c)** Average EPSPs under different pharmacological conditions shown in **(b)**. The presynaptic APs are shown at the top. The PPR values of different traces are indicated on the right.
